# Tensor Decomposition and Principal Component Analysis-Based Unsupervised Feature Extraction Outperforms State-of-the-Art Methods When Applied to Histone Modification Profiles

**DOI:** 10.1101/2022.04.29.490081

**Authors:** Sanjiban Sekhar Roy, Y-h. Taguchi

## Abstract

Identification of histone modification from datasets that contain high-throughput sequencing data is difficult. Although multiple methods have been developed to identify histone modification, most of these methods are not specific for histone modification but are general methods that aim to identify protein binding to the genome. In this study, tensor decomposition (TD) and principal component analysis (PCA)-based unsupervised feature extraction with optimized standard deviation were successfully applied to gene expression and DNA methylation. The proposed method was used to identify histone modification. Histone modification along the genome is binned within the region of length *L*. Considering principal components (PCs) or singular value vectors (SVVs) that TD or PCA attributes to samples, we can select PCs or SVVs attributed to regions. The selected PCs and SVVs further attribute *P*-values to regions, and adjusted P-values are used to select regions. The proposed method identified various histone modifications successfully and outperformed various state-of-the-art methods. This method is expected to serve as a *de facto* standard method to identify histone modification.

## 1 Introduction

Identification of histone modification [1] from high-throughput sequencing (HTS) datasets is important for the following reasons.

- Histone modification contributes to various functional genomic features, including transcription [2] and alteration of chromatin structures [3].
- In contrast to variable gene expression, histone modification is more stable [4]; thus, it may be used to characterize the state of the genome.
- In contrast to DNA methylation that has only two states, methylated or not, histones are modified in various ways. Thus, histone modification is related to a more detailed functionality of the genome [5].
- Since histone modification may activate or suppress transcription, combinations of various histone modifications may have more complicated transcriptional roles [6].

In spite of the importance of histone modification, gene expression and DNA methylation have instead been the focus of genomic studies for the following reasons.

- To identify histone modification throughout the genome, detection of antibody binding to the entire genome is required [7]. In contrast, gene expression can be detected even if only transcription start sites are sequenced.
- In contrast to gene expression that can be measured even when only exons are considered or DNA methylation that is meaningful even if only promoter regions are considered, specific regions of the genome cannot be used due to limited knowledge on the position-specific functionality of histone modification [8].
- Because of the above two reasons, HTS for histone modification requires more depth, which is both time-consuming and expensive [9].
- Similarly, the number of datasets and computational resources required to identify histone modification is greater than that required to measure gene expression and DNA methylation [10].

Possibly because of the unpopularity of histone modification identification, fewer methods specific to histone modification have been developed. Because histone modification is often measured using chromatin immunoprecipitation sequencing (ChIP-seq) technology to identify protein binding sites on DNA [9], so-called peak call programs [11] developed to process protein binding to DNA via ChIP-seq have often been used to process HTS datasets used for histone modification. However, since histone modifications are hardly regarded to be distributed over the genome by forming peaks, these peak calling algorithms are not guaranteed to identify histone modification. For example, most methods employ binomial distributions to identify the amount of histone modification over the genome [12], but we are unsure if these methods are suitable. A more flexible method that does not assume a specific distribution of histone modification over the whole genome is required.

To fulfill this requirement, the method of tensor decomposition (TD) and principal component analysis (PCA)- based unsupervised feature extraction (FE) with optimized standard deviation (SD) that were successfully applied to gene expression [13] and DNA methylation [14] were tested to determine if histone modification could also be identified. In contrast to two previous studies [13], [14] where PC or singular value vectors (SVVs) obey the Gaussian distribution which is assumed in the null hypothesis after SD optimization, in our method PC and SVVs follow a mixed Gaussian distribution instead. Nevertheless, empirically, SD optimization allows the correct identification of histone modification to some extent, even when compared to various state-of-the-art methods.

## 2 Materials and Methods

### 2.1 Histone modification profiles

The following histone modification profiles have been used to evaluate the performance of the proposed method. All profiles were retrieved from the Gene Expression Omnibus (GEO) database [15].

#### 2.1.1 GSE24850

This study contained eleven H3K9me3 ChIP-seq mouse nucleus accumbens experiments comprising six controls, three saline-treated samples, and two cocaine samples [16]. Among them, five controls and five treated samples were employed. Ten bed files that were provided as a Supplementary file in GEO were downloaded. One control samples with a GEO identification (ID) of GSM612984 was discarded and the other ten samples were used.

#### 2.1.2 GSE159075

This study contained various histone modification ChIP-seq experiments using human umbilical vein endothelial cell lines [17]. Among them three H3K4me3 ChIP-seq, three H3K27me3 ChIP-seq, and three H3K27ac ChIP-seq, as well as one input ChIP-seq profile were employed. The corresponding 10 bigWig files were downloaded from the GEO Supplementary file.

#### 2.1.3 GSE74055

This study contained various histone modification ChIP-seq experiments using mouse E14 or DKO cell lines [18]. Among them, 16 H3K4me1 ChIP-seq profiles and the corresponding 16 input profiles (in total 32 profiles) were downloaded in the bigWig format provided in the GEO Supplementary file.

#### 2.1.4 GSE124690

This study has comprised various histone modification CUT&Tag experiments using human H1 or K562 cell lines [19]. Among them, six bulk H3K4me1 profiles (GSM3536499_H1_K4me1_Rep1.bed.gz, GSM3536499_H1_K4me1_Rep2.bed.gz, GSM3536516_K562_K4me1_Rep1.bed.gz, GSM3536516_K562_K4me1_Rep2.bed.gz, GSM3680223_K562_H3K4me1_Abcam_8895.bed.gz, GSM3680224_K562_H3K4me1_ActMot_39113.bed.gz), four bulk H3K3me3 profiles (GSM3536501_H1_K4me3_Rep1.bed.gz, GSM3536501_H1_K4me3_Rep2.bed.gz, GSM3536518_K562_K4me3_Rep1.bed.gz, GSM3536518_K562_K4me3_Rep2.bed.gz), and four bulk H3K27ac profiles (GSM3536497_H1_K27ac_Rep1.bed.gz, GSM3536497_H1_K27ac_Rep2.bed.gz, GSM3536514_K562_K27ac_Rep1.bed.gz, GSM3536514_K562_K27ac_Rep2.bed.gz) were downloaded from the GEO Supplementary file.

#### 2.1.5 GSE188173

This study contained nine ChIP-seq H3K27ac profiles (with one control and one treated with SPT) using patient–derived xenografts of human castration-resistant prostate cancer (18 profiles in total). The corresponding 18 bigWig files were downloaded from the GEO Supplementary file.

#### 2.1.6 GSE159022

This study comprised four H3K4me3 ChIP-seq profiles, four H3K27me3 ChIP-seq profiles, and four H4K16ac ChIP-seq profiles using mouse progenitor cells (two wild type (WT) and two neurofibromin knockouts) [20]. Among them, four H3K27me3 profiles were used and four bigWig files were downloaded from the GEO Supplementary file.

#### 2.1.7 GSE168971

This study contained H3K27ac and H3K9ac ChIP-seq profiles taken from various experimental conditions [21]. Among them, six H3K9ac profiles using C3H-WT mouse liver and the two corresponding inputs were used. The corresponding eight bigWig files were downloaded from the GEO Supplementary file.

#### 2.1.8 GSE159411

This study comprised various ChIP-seq profiles [22]. Among them, four H3K36me3 ChIP-seq profiles (two hiPSC cardiomyocyte and two WT hiPSC) were used. The four corresponding bigWig files were downloaded from the GEO Supplementary file.

#### 2.1.9 GSE181596

This study consisted of four H3K27me3 ChIP-seq profiles (two controls and two treatments) and four H3K4me3 ChIP-seq profiles (two controls and two treatments) in addition to two input profiles that used cells as odontoblasts (treatment was siRNA: si-IKBz) [23]. Among them, four H3K27me3 ChIP-seq profiles were downloaded from the GEO Supplementary file.

### 2.2 Histone modification profile pre-processing

To apply the proposed method to histone modification profiles, individual histone modification profiles must share region index *i*. Therefore, the amount of histone modification is averaged within shared regions that is generated by dividing the whole human or mouse genome into equal length, *L,* intervals in each chromosome. *L* = 1000 is used for all histone profiles except H3K9me3, for which *L* = 25000 is used.

There are two kinds of bed formats for histone modification. One is coarse grained histone modification, where histone modification is averaged within intervals; the other is the genomic loci where histone modification occurs. For the former, coarse grained values are further averaged with the region of length *L*; for the latter, the regions overlapping the regions of length *L* are counted.

### 2.3 PCA-based unsupervised FE with optimized SD

In our work the PCA-based unsupervised FE with optimized SD method was used. In the literature, few proposed methods have dealt with PCA-based unsupervised feature extraction; such methods could be found in previous studies [?], [14], It is briefly described below.

Suppose histone modification profiles are formatted as a matrix *x_ij_* ∈ ℓ^*N*×*M*^ that represents histone modification of ith regions in *j*th samples. Here, we assume that *x_ij_* is normalized as

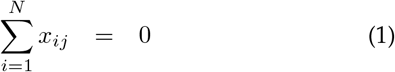

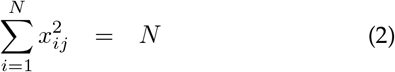

Applying PCA to *x_ij_* should obtain

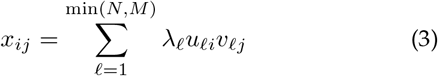

where

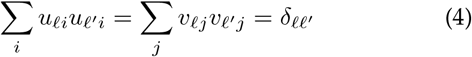

To get *u_ℓ_i__*, the eigenvalues and vector of 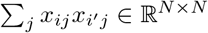 must be obtained as

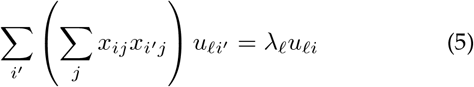

and *v_ℓj_* can be obtained as

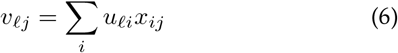

To select regions of interest, *i*, first, the *v_ℓj_* associated with the desired property is identified. In this study, the property of interest is histone modification that is independent of samples, that is, *v_ℓj_* should be independent of *j*. After identifying *ℓ* of interest, an optimal SD of *u_ℓi_* is obtained such that *u_ℓi_* obeys the Gaussian distribution (null hypothesis) as much as possible.

SD optimization can be done as follows. First, *P*-values are attributed to the *i*th region as

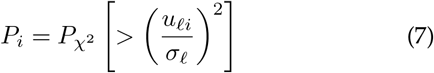

where 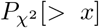 is the cumulative *χ*^2^ distribution where the argument is larger than x and *σ_ℓ_* is the SD. Then, the histogram *h_s_* of *P_i_* is computed, which is the number of is that satisfy

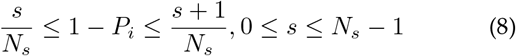

*h_s_* is expected to have constant values for *s* ≤ *s*_0_, with some *s*_0_, and a sharp peak exists at *k*_0_ < *k*. In this case, is included in *h_s_, s* > *s*_0_ are selected based on association with significant histone modification. Next, *P_i_*s are recomputed with optimized SD and the obtained *P_i_*s are corrected using BH criterion [24] and is with adjusted *P_i_*s less than 0.01 are selected.

### 2.4 TD-based unsupervised FE with optimized SD

Below we show how PCA is replaced with TD. Suppose that histone modification is formatted as a tensor *x_ijk_* ∈ ℓ^*N*×*M*×*K*^ in the *i*th region of the *j*th sample under the *k*th condition and TD is obtained as

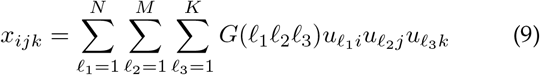

where *G* ∈ ℓ^*N*×*M*×*K*^ and

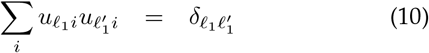

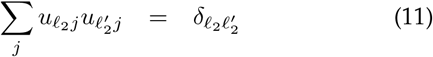

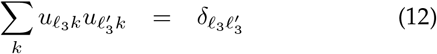

with higher order singular value decomposition (HOSVD [24]). After identifying *ℓ*_2_ and *ℓ*_3_ of interest by investigating the dependence of *u*_*ℓ*_2_*j*_ and *u*_*ℓ*_3_*k*_ on *j* and *k*, the *u*_*ℓ*_1_*i*_ associated with absolutely the largest *G* is selected using the selected *ℓ*_2_ and *ℓ*_3_. The following procedure is the same as above.

### 2.5 Performance evaluation by Enrichr

After the regions were selected, Entrez gene IDs included in these regions were listed and were converted to gene symbols using the gene ID converter implemented in DAVID [25], [26]. The obtained list of gene symbols was uploaded to Enrichr [27]. The primary “Epigenomics Roadmap HM ChIP-seq” category was employed to evaluate the performance and the “ENCODE Histone Modifications 2015” category was additionally used for the evaluation if no significant results were obtained in the “Epigenomics Roadmap HM ChIP-seq” category using the proposed method.

### 2.6 Methods for comparison

The methods used for comparison in the proposed method were selected based on the following criteria.

- The method must accept bed or bigWig file formats as input since the proposed method cannot accept the sam/bam format as input, which is used by the most popular methods.
- The method must be standalone (not web-based).
- The method must be performed in the Linux platform.
- The method must be implemented as free (open) software.

The dataset tested was GSE24850 (H3K9me3).

#### 2.6.1 MOSAiCS

MOSAiCS [28] was implemented as a ioconductor package [29]. Version 2.32.0 was installed in R [30] and applied to GSE24850. MOSAiCS provides biologically motivated statistical models for reads that arise under both non-enrichment (background) and enrichment (signal). Furthermore, MOSAiCS builds a parametric background model that takes into account biases such as GC content and mappability that are inherent to ChIP-seq data. The MOSAiCS model does not assume punctuated or broad peak structures but instead quantifies whether the ChIP reads show enrichment compared to the background reads for every genomic interval (e.g., bin) of user defined size in the genome.

#### 2.6.2 DFilter

DFilter [31] was implemented as a Linux command-line program. Ver. 1.6 was downloaded from the website https://reggenlab.github.io/DFilter/. DFilter takes as input a set of sequence tags mapped to a reference genome. Based on the genomic distribution of tags, the algorithm classifies individual n-base-pair bins as positive (signal) or negative (noise) regions. DFilter implements linear finite-impulse-response detection, that is, a windowed linear filter h of user-specified width, followed by the standard thresholding step.

#### 2.6.3 F-Seq2

F-Seq2 [32] was implemented as a Linux command line program. It was downloaded from Github https://github.com/Boyle-Lab/F-Seq2. F-Seq2 employed Gaussian kernel density function to quantify the amount of protein binding. The total control read count was linearly scaled to be equal to the total treatment read count at the individual chromosome level as the ratios of total reads fluctuated between different chromosomes.

#### 2.6.4 HOMER

HOMER [33] was implemented as a Linux command line program. The latest version, HOMER 4.11, which was released on October 24, 2019, was downloaded from the Web site http://homer.ucsd.edu/homer/. For each ChIP-Seq experiment, ChIP-enriched regions (peaks) were found by first identifying significant clusters of ChIP-Seq tags and then filtering these clusters for those significantly enriched relative to background sequencing and local ChIP-Seq signal.

#### 2.6.5 RSEG

RSEG [34] was implemented as a Linux command line program. The latest version, 0.4.9, was downloaded from the web site http://smithlabresearch.org/software/rseg/. Negative binomial distribution is assumed to quantify the amount of protein binding between control and treated samples using the NBDiff distribution, which is the discrete distribution of the difference between two independent negative binomial random variables.

### 2.7 Gaussian Mixture Analysis

Gaussian mixture analysis was performed using `mclust` package [35] in R [30]. The Mclust function was applied to *u*_1*i*_ for GSE24850.

## 3 Results

A full list of genes and the enrichment analysis is in the Supplementary materials.

### 3.1 GSE24850

#### 3.1.1 PCA-based unsupervised FE with optimized SD

The proposed method was applied to H3K9me3 modifications in the GSE24850 dataset. PCA was applied to three saline-treated samples and two cocaine samples (five samples in total). The first PC loading, *v*_1*j*_, attributed to samples was independent of *j* attributed to samples. Then, the corresponding first PC score, *u*_1*i*_, was used for gene selection. SD was optimized and the SD = 0.04264199. The histogram of 1 – *P_i_* appeared to be a combination of two Gaussian distributions as opposed to a single Gaussian distribution (Fig. 2). A small number of regions (1302) were selected among a total of 106,204 regions (i.e., only a few percentages) and the associated 894 Entrez gene IDs were identified. The gene IDs were converted to 641 gene symbols, which were uploaded to Enrichr. Table 1 shows the enriched histone modification associated with adjusted *P* values less than 0.05. All were H3K9me3 modifications, suggesting that the proposed method was very successful. In addition, five out of ten were brain related, which was coincident with the fact that GSE24850 comprised experiments using the nucleus accumbens, strengthening the success of the proposed method.

**Fig. 1.**
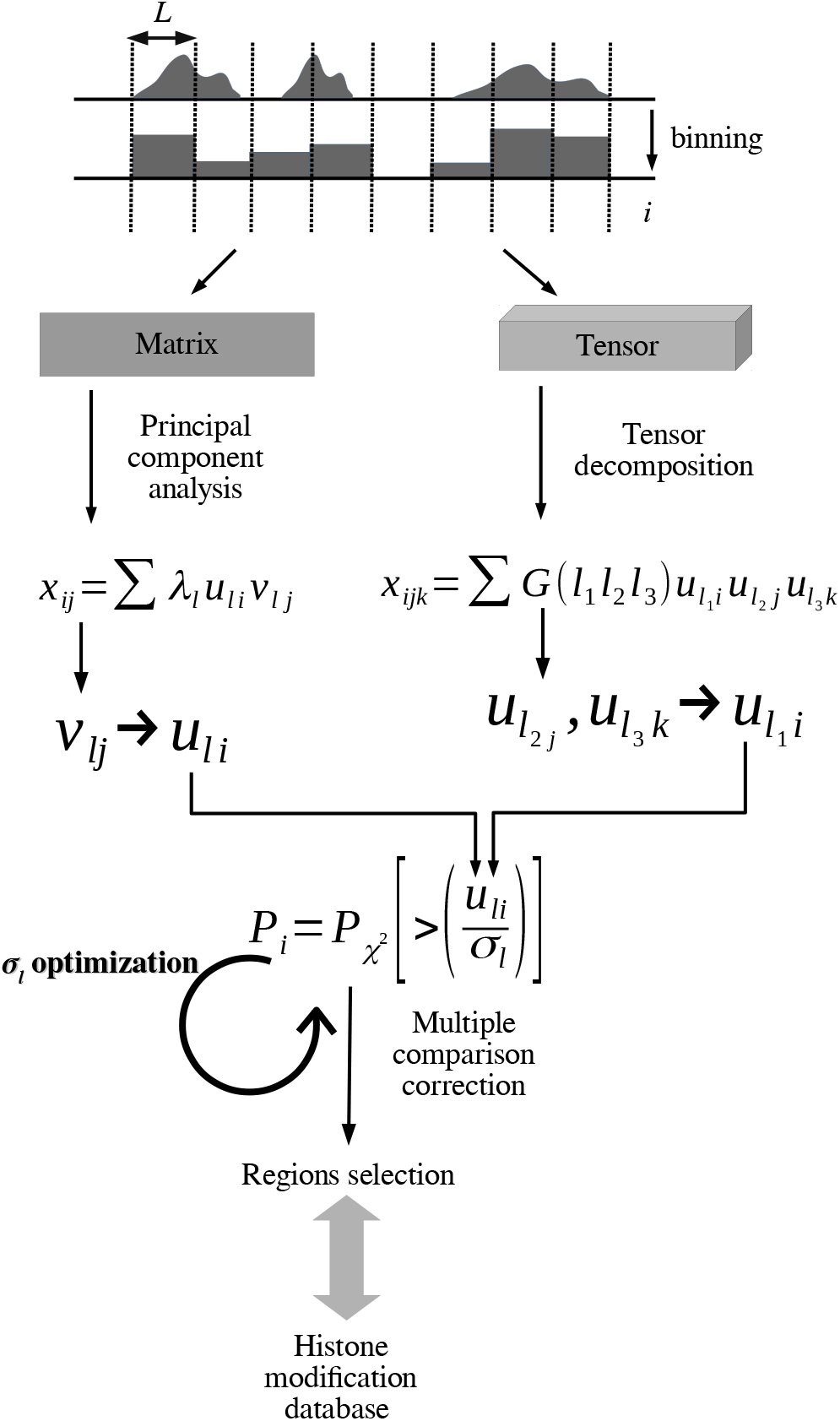
Schematic diagram for processing histone modification data. Histone modification is binned within the region of length *L*. Binned profiles are integrated as matrices or tensors to which PCA or TD is applied. Obtained *v_ℓj_* or *u*_*ℓ*_2_*j*_,*u*_*ℓ*_3_*k*_ attributed to samples is used to identify which *u_ℓ_i__* or *u*_*ℓ*_1_*i*_ is used to attribute the *P*-value, *P_i_*, to the ith region. *P_i_*s are corrected by BH criterion and regions associated with adjusted *P*-values less than 0.01 are selected. Enrichment of histone modification in databases are investigated toward the selected regions.

**Fig. 2.**
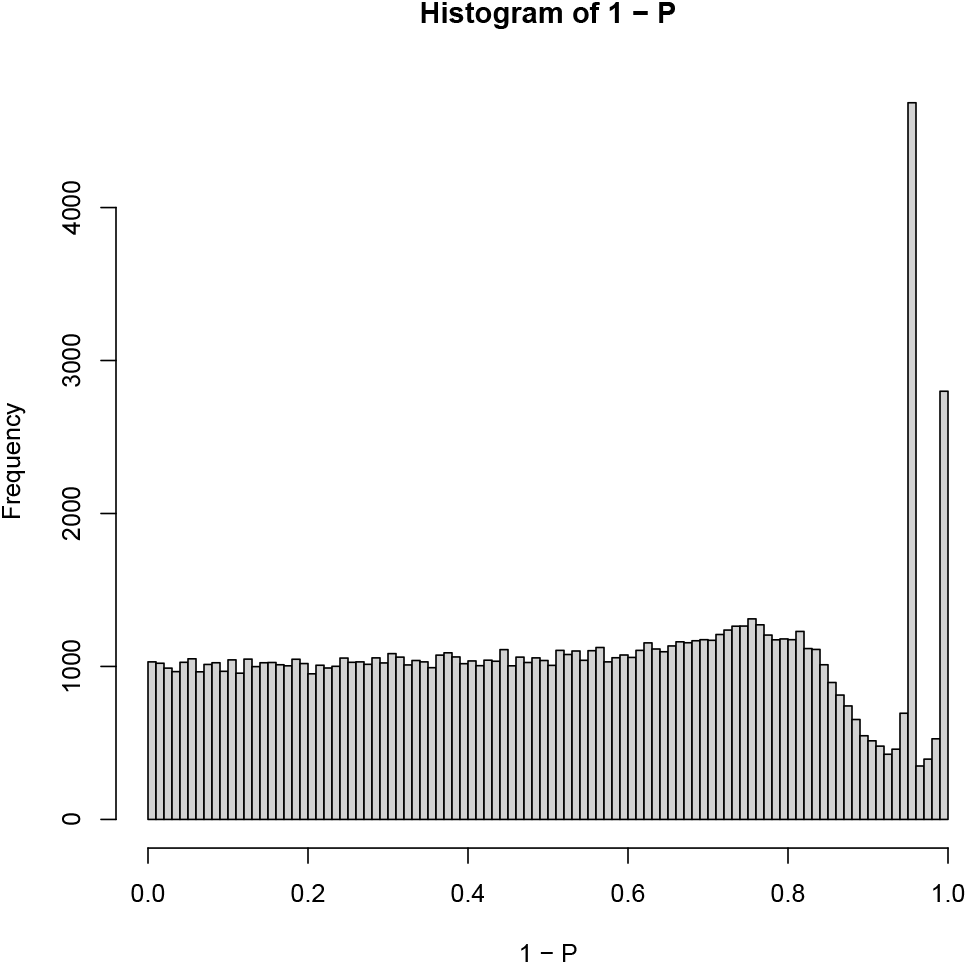
Histogram of 1 - *P_i_* computed using the proposed method for the GSE24850 dataset (H3K9me3)

**TABLE 1.** Enriched histone modification in the “Epigenomics Roadmap HM ChIP-seq” category of Enrichr for the GSE24850 dataset (H3K9me3)

| Term | Overlap | P-value | Adjusted P-value |
| --- | --- | --- | --- |
| H3K9me3 Brain Mid Frontal Lobe | 22/217 | $2.02 \times 10^{-6}$ | $7.45 \times 10^{-4}$ |
| H3K9me3 Brain Inferior Temporal Lobe | 18/170 | $9.31 \times 10^{-6}$ | $1.72 \times 10^{-3}$ |
| H3K9me3 CD4 Naive Primary Cells | 10/64 | $3.39 \times 10^{-5}$ | $4.17 \times 10^{-3}$ |
| H3K9me3 Brain Anterior Caudate | 15/141 | $4.82 \times 10^{-5}$ | $4.45 \times 10^{-3}$ |
| H3K9me3 IMR90 | 45/814 | $2.82 \times 10^{-4}$ | $2.08 \times 10^{-2}$ |
| H3K9me3 Brain Hippocampus Middle | 15/170 | $3.90 \times 10^{-4}$ | $2.40 \times 10^{-2}$ |
| H3K9me3 Stomach Smooth Muscle | 17/217 | $6.56 \times 10^{-4}$ | $3.33 \times 10^{-2}$ |
| H3K9me3 Colon Smooth Muscle | 10/92 | $7.24 \times 10^{-4}$ | $3.33 \times 10^{-2}$ |
| H3K9me3 CD8 Naive Primary Cells | 11/110 | $8.11 \times 10^{-4}$ | $3.33 \times 10^{-2}$ |
| H3K9me3 Brain Cingulate Gyrus | 11/115 | $1.17 \times 10^{-3}$ | $4.33 \times 10^{-2}$ |

#### 3.1.2 Comparisons with state-of-the-art methods

Although the proposed method successfully achieved a significant performance, if other state-of-the-art methods achieve a similar performance, the importance of the proposed method drastically decreases. To reject this possibility, various state-of-the-art methods were used to analyze the GSE24850 dataset. No state-of-the-art method had a comparative performance.

##### 3.1.2.1 MOSAiCS

The first state-of-the-art method tested was MOSAiCS. Since MOSAiCS only accepts a pair of ChIP-seq and input datasets (i.e., a control), MOSAiCS was repeatedly applied to five pairs of ChIP-seq data (i.e., three saline and two cocaine samples) and one input profile. MOSAiCS identified 4367, 3648, 2096, 1985, and 5566 peak regions and 1833, 1599, 1136, 1018, and 2223 associated Entrez gene IDs. Finally, 994, 851, 567, 532, and 1184 gene symbols were identified. Next, these five sets of genes were uploaded to Enrichr one by one and the number of histone modifications that were significantly enriched in the “Epigenomics Roadmap HM ChIP-seq” category of Enrichr was determined. Table 2 shows the performance of MOSAiCS, which identifies at most only three significantly enriched histone modifications, at most two of which are H3K9me3 modifications. Since the proposed method identified as large as ten enriched histone modification all of which are H3K9me3 modifications, the proposed method outperforms MOSAiCS.

**TABLE 2.** The number of regions/peaks, Entrez gene IDs, and gene symbols selected by various methods and that of the H3K9me3-associated enriched histone modification in the “Epigenomics Roadmap HM ChIP-seq” category of Enrichr with adjusted *P*-values less than 0.05 for the GSE24850 dataset (H3K9me3).

| Methods | Pair no. | Regions/peaks | Entrez genes | Gene symbols | Enriched histone modification | H3K9me3 |
| --- | --- | --- | --- | --- | --- | --- |
| Proposed method | — | 1302 | 894 | 641 | 10 | 10 |
| MOSAICS | 1 | 4367 | 1833 | 994 | 3 | 1 |
|  | 2 | 3648 | 1599 | 851 | 0 | 0 |
|  | 3 | 2096 | 1136 | 567 | 0 | 0 |
|  | 4 | 1985 | 1018 | 532 | 2 | 2 |
|  | 5 | 5556 | 2223 | 1184 | 2 | 0 |
| DFilter | 1 | 25080 | 6286 | 2621 | 1 | 0 |
|  | 2 | 22863 | 5721 | 2524 | 1 | 0 |
|  | 3 | 21371 | 5470 | 2499 | 1 | 0 |
|  | 4 | 23811 | 5987 | 2631 | 1 | 0 |
|  | 5 | 23369 | 5902 | 2544 | 1 | 0 |
| F-Seq2 | — | — | — | — | — | — |
| HOMER | — | 114727 | 6771 | 6747 | 1 | 0 |
| RSEG | — | — | — | — | — | — |

##### 3.1.2.2 DFilter

The next state-of-the-art method tested was DFilter, which was also applied to the five pairs of data. DFilter identified 25080, 22863, 21371, 23811, and 23369 peak regions and 6286, 5721, 5407, 5987, and 5903 Entrez gene IDs, of which 2621, 2524, 2499, 2631, and 2544 gene symbols were associated with each of the five pairs. These sets of gene symbols were uploaded to Enrichr and no enriched H3K9me3 modifications were observed (Table 2). Thus, the proposed method outperforms DFilter.

##### 3.1.2.3 F-Seq2

The F-Seq2 method could not be performed with the available computer memory. Therefore, we could not compare the performance between F-Seq2 and the proposed method.

##### 3.1.2.4 HOMER

The next state-of-the-art method was HOMER, which has been used in the ENCODE project and has been used for recent reports [36]. HOMER has the capability to compare five input profiles with five H3K9me3 profiles directly. Using this capability, HOMER identified 114727 peak regions, 6771 associated Entrez genes, and 6747 gene symbols which were uploaded to Enrichr. Unfortunately, Enrichr did not identify enriched regions of H3K9me3 modifications (Table 2).

##### 3.1.2.5 RSEG

The last comparison was with the state-of-the-art method RSEG, which resulted in a core dump, although no compile errors were detected. The RSEG demo file also could not be treated without errors; it was too old to be used on the present platform. Although it has been cited many times as one of the popular tools to process ChIP-seq datasets, it has not been tested in recent publications.

### 3.2 Histone modification other than H3K9me3

Although the superiority of the proposed method has been demonstrated using H3K9me3 profiles, one of five “core histone marks” proposed by the Roadmap Epigenomics Consortium [37] (H3K4me1/H3K27ac, H3K4me3, H3K36me3, H3K27me3, and H3K9me3), there is a possibility that the proposed method is not effective with other histone modifications. To reject this possibility, the proposed method was tested on various other histone modifications.

#### 3.2.1 GSE159075

Regions, *is*, with non-zero missing values among three samples were discarded in advance. PCA was applied to three samples with the same histone modification (H3K4me3, H3K27me3, or H3K27ac) and *v*_1*j*_ was always associated with the independence of *j* (biological replicates). Next, the corresponding first PC score, *u*_1*i*_, was used for gene selection. SDs were optimized and the SD = 0.19218492 (H3K4me3), 0.8650455 (H3K27me3), and 0.3911769 (H3K27ac). The histogram of 1 – *P_i_* seemed to obey a combination of two Gaussian distributions, and not a single Gaussian distribution (Fig. 3). As a result, 34538, 62141, and 61306 regions were selected, and 13692, 5217, and 11604 Entrez gene IDs were identified. The 13671, 5208, and 11590 gene symbols associated with the gene IDs were identified and uploaded to Enrichr. Table 3 shows the performance evaluation by Enrichr and demonstrates the success of the proposed method.

**Fig. 3.**
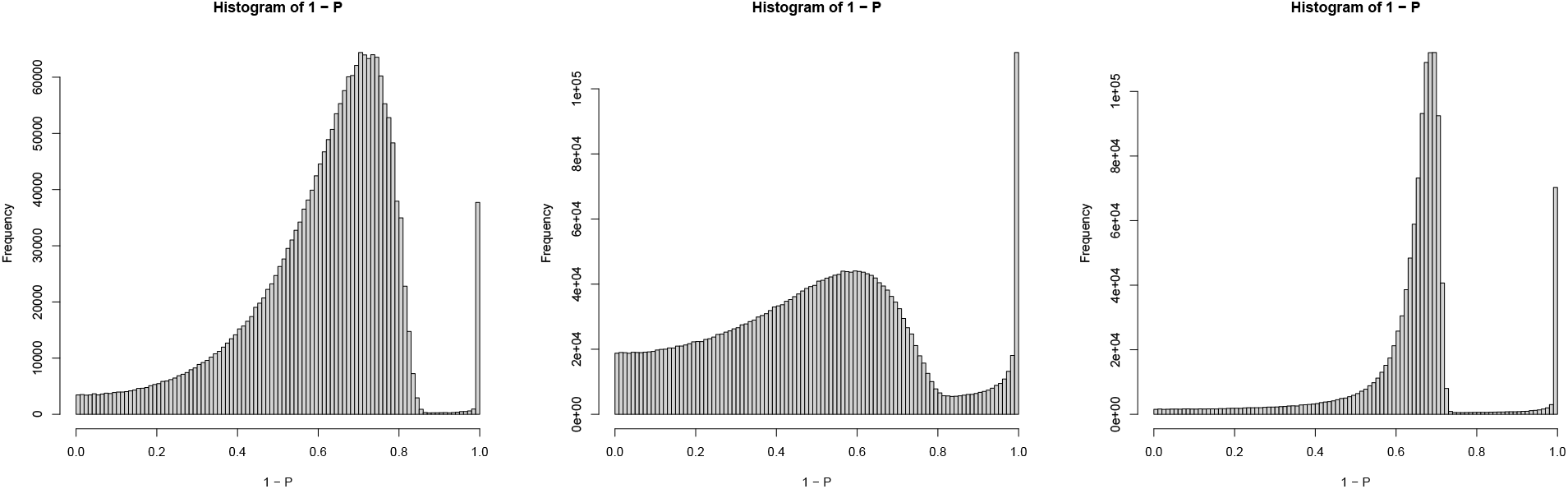
Histogram of 1 - *P_i_* computed by the proposed method using the GSE159075 dataset (from left to right, H3K4me3, H3K27me3, and H3K27ac)

**TABLE 3.**
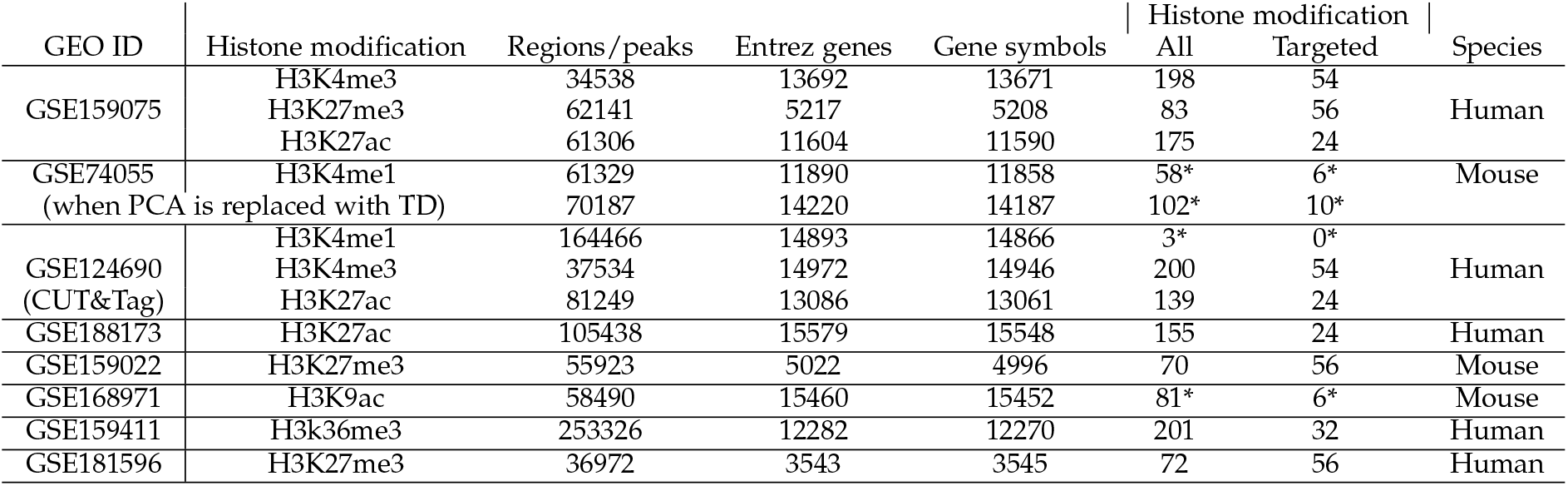
The number of regions/peaks, Entrez gene IDs, and gene symbols selected by various methods and that of associated enriched histone modification (all and targeted) in the “Epigenomics Roadmap HM ChIP-seq” category of Enrichr with adjusted P-values less than 0.05 for other profiles than GSE24850 shown in Table 1. *:ENCODE Histone Modifications 2015

#### 3.2.2 GSE74055

PCA was applied to 16 H3K4me1 modifications divided by the corresponding input (Regions, *i*s, for which input samples were zero, were discarded in advance). The *u*_1*i*_ could not provide reasonable results, so *u*_2*i*_ was used for gene selection. SDs were optimized and the SD = 0.1873585. Figure 4 shows the histogram of 1 – *P_i_* with optimized SD. Next, 61329 regions, 11890 Entrez gene IDs, and 11858 gene symbols were identified. The gene symbols were uploaded to Enrichr. Table 3 shows the performance evaluation by Enrichr. Its superiority was reduced since no enriched H3K4me1 modifications were identified for the “Epigenomics Roadmap HM ChIP-seq” category. The other category “ENCODE Histone Modifications 2015”, which had more enriched histone modifications, needs to be consulted. Regardless, non-zero enriched H3K4me1 profiles could be detected in Enrichr.

**Fig. 4.**
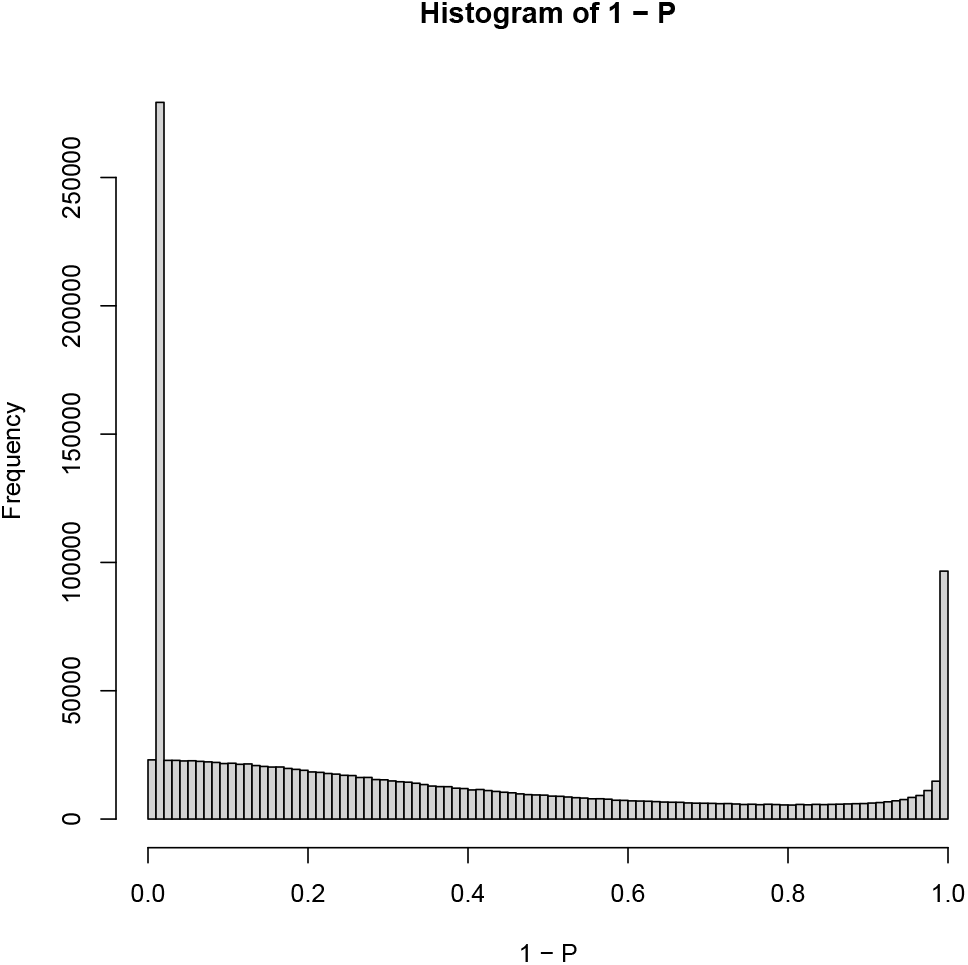
Histogram of 1 - *P_i_* computed by the proposed method using the GSE74055 dataset (H3K4me1)

#### 3.2.3 GSE124690

PCA was applied to six H3K4me1, four H3K4me3, and four H3K27ac ChIP-seq profiles separately and *v*_1*j*_ was always associated with the independence of *j* (biological replicates). However, *v*_2*j*_ was used for H3K4me1 modifications as *u*_1*i*_ could not provide good results. Next, the corresponding first PC score, *u*_1*i*_, was used for gene selection for H3K4me3 and H3K27ac modifications and the second PC score, *u*_2*i*_, was used for gene selection for H3K4me1 modifications. SDs were optimized and the SD = 0.5468109 (H3K4me1), 0.5600861 (H3K4me3), and 0.4509331 (H3K27ac). The histogram of 1 – *P_i_* seemed to obey a combination of two Gaussian distributions and not a single Gaussian distribution (Fig. 5). A total of 164466, 37534, and 81249 regions, 14893, 14972, and 13061 Entrez gene IDs, and 14866, 14946, and 13061 gene symbols (2nd PC was used for H3K4me1) were identified. The gene symbols were uploaded to Enrichr. Table 3 shows the performance evaluation by Enrichr. Again, its superiority toward H3K4me1 modifications was reduced since no enriched H3K4me1 modifications were identified for the “Epigenomics Roadmap HM ChIP-seq” or “ENCODE Histone Modifications 2015” categories. For the other two histone modifications, Enrichr was very successful.

**Fig. 5.**
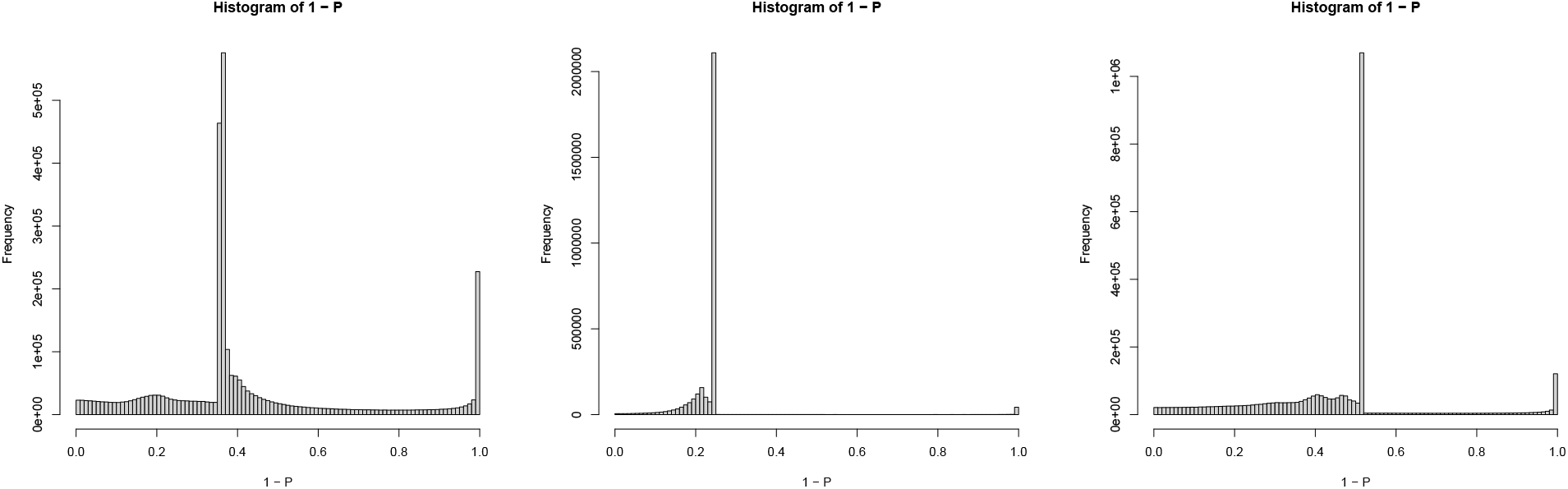
Histogram of 1 - *P_i_* computed by the proposed method using the GSE124690 dataset (From left to right, H3K4me1, H3K4me3, and H3K27ac)

#### 3.2.4 GSE188173

PCA was applied to 16 H3K27ac profiles and *v*_1*j*_ was always associated with the independence of *j* (biological replicates). Next, the corresponding first PC score, *u*_1*i*_, was used for gene selection. SDs were optimized, and the SD = 0.5319398. Figure 6 shows the histogram of 1 – *P_i_* with optimized SD. A total of 105438 regions, 15579 Entrez gene IDs, and 15548 gene symbols were identified that were uploaded to Enrichr. Table 3 shows the performance evaluation by Enrichr, which demonstrates the success of the proposed method.

**Fig. 6.**
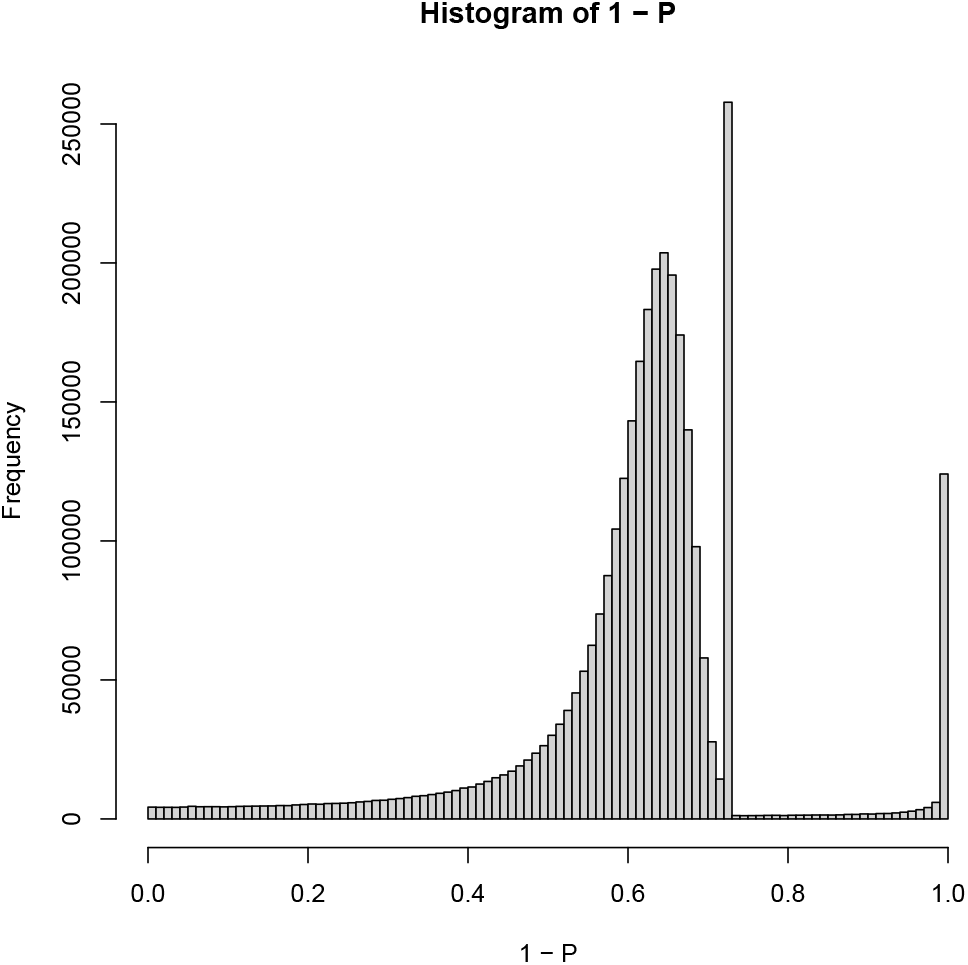
Histogram of 1 - *P_i_* computed by the proposed method using the GSE188173 dataset (H3K27ac)

#### 3.2.5 GSE159022

PCA was applied to H3K27me3 ChIP-seq profiles and *v*_1*j*_ was always associated with the independence of *j* (biological replicates). Then, the corresponding first PC score, *u*_1*i*_, was used for gene selection. SDs were optimized and the SD = 0.06544008. Figure 7 shows the histogram of 1 — *P_i_* with optimized SD. A total of 55923 regions, 5022 Entrez gene IDs, and 4996 gene symbols were identified. The gene symbols were uploaded to Enrichr. Table 3 shows the performance evaluation by Enrichr, which demonstrates the success of the proposed method.

**Fig. 7.**
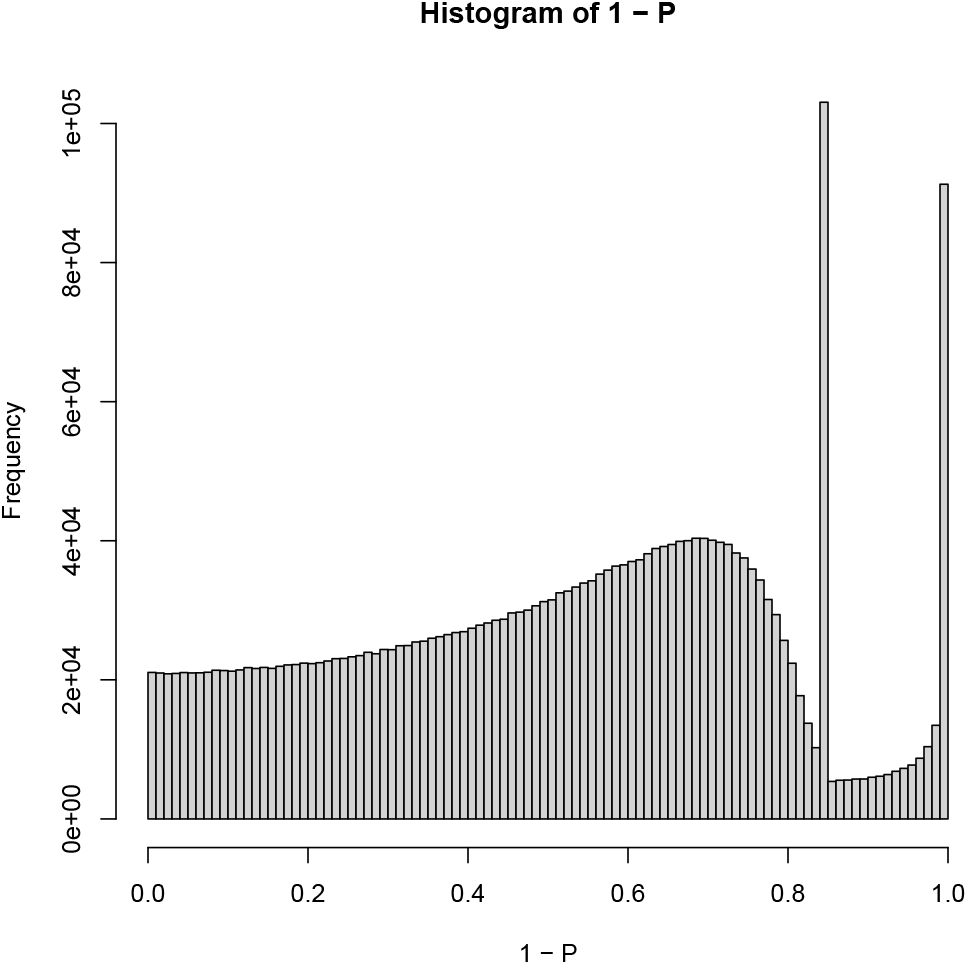
Histogram of 1 - *P_i_* computed by the proposed method using the GSE159022 dataset (H3K27me3)

#### 3.2.6 GSE168971

PCA was applied to six H3K9ac ChIP-seq profiles and two corresponding inputs. *v*_1*j*_ was distinct between treated and control samples. Next, the corresponding first PC score, *u*_1*i*_, was used for gene selection. SDs were optimized and the SD = 0.3697877. Figure 8 shows the histogram of 1 – *P_i_* with optimized SD. A total of 58490 regions, 15460 Entrez gene IDs, and 15452 gene symbols were identified. The gene symbols were uploaded to Enrichr. Table 3 shows the performance evaluation by Enrichr. Its performance for H3K9ac modifications was reduced since the “ENCODE Histone Modifications 2015” category had to be consulted and it only identified six enriched H3K9ac profiles in the “ENCODE Histone Modifications 2015” category.

**Fig. 8.**
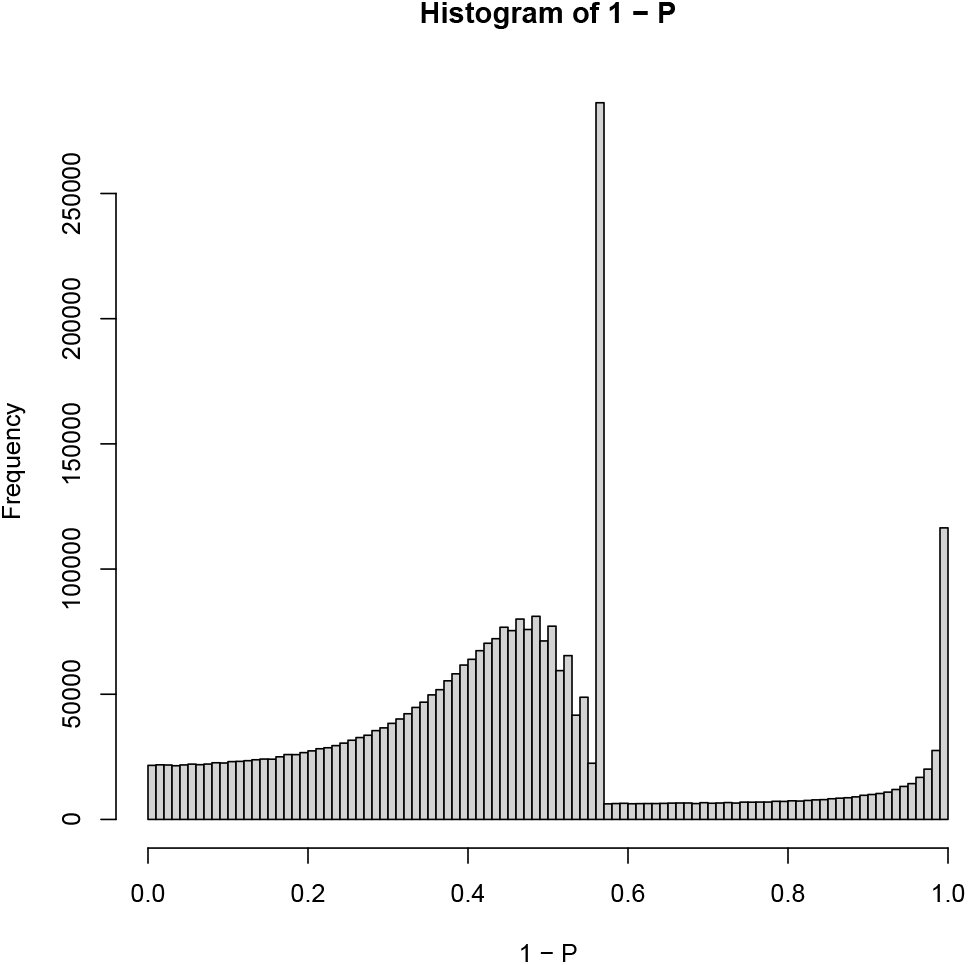
Histogram of 1 - *P_i_* computed by the proposed method using the GSE168971 dataset (H3K9ac)

#### 3.2.7 GSE159411

PCA was applied to four H3k36me3 ChIP-seq profiles. *v*_1*j*_ was always associated with the independence of *j* (biological replicates). Then, the corresponding first PC score, *u*_1*i*_, was used for gene selection. SDs were optimized and the SD = 0.6005592. Figure 9 shows the histogram of 1 – *P_i_* with optimized SD. A total of 25,3326 regions, 12282 Entrez gene IDs, and 12270 gene symbols were identified. The gene symbols were uploaded to Enrichr. Table 3 shows the Enrichr performance evaluation, which demonstrates the success of the proposed method.

**Fig. 9.**
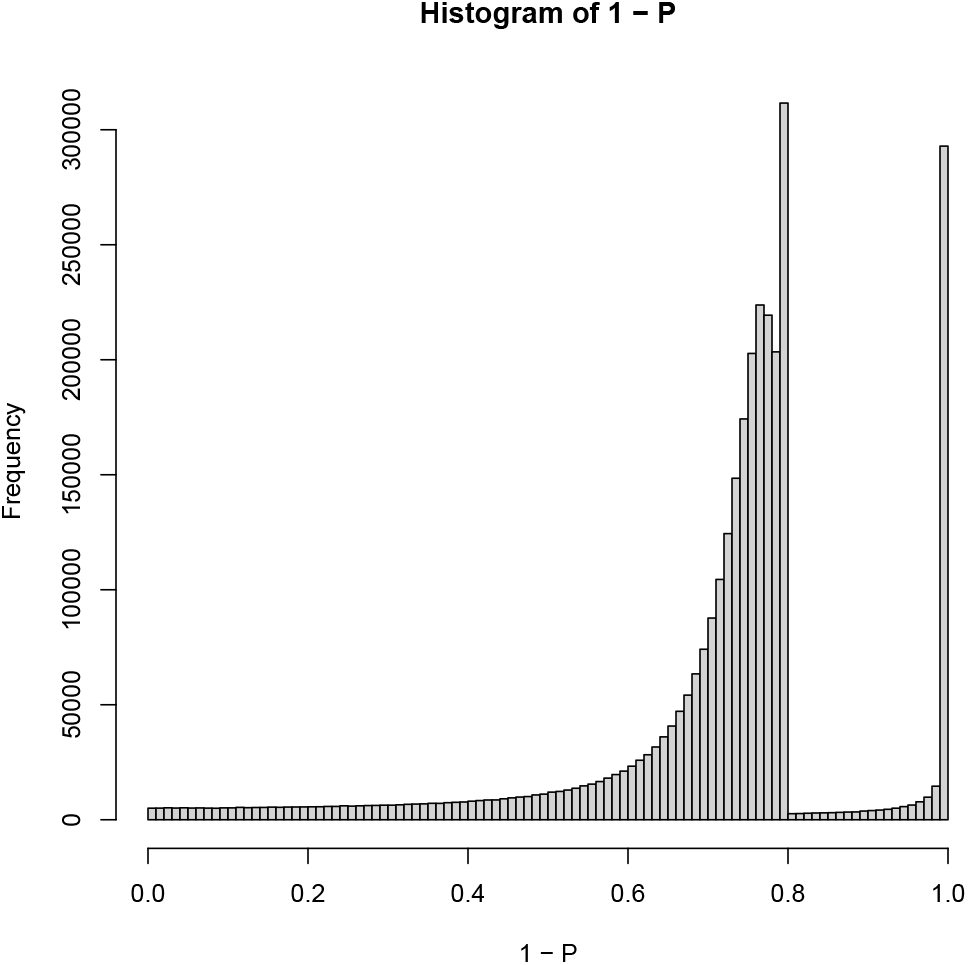
Histogram of 1 - *P_i_* computed by the proposed method using the GSE159411 dataset (H3K36me3)

#### 3.2.8 GSE181596

PCA was applied to four H3K27me3 ChIP-seq profiles. *v*_1*j*_ was always associated with the independence of *j* (biological replicates). Next, the corresponding first PC score, *u*_1*i*_, was used for gene selection. SDs were optimized, and the SD = 1.855494. Figure 10 shows the histogram of 1 – *P_i_* with optimized SD. A total of 36972 regions, 3543 Entrez gene IDs, and 3545 gene symbols were identified. The gene symbols were uploaded to Enrichr. Table 3 shows the Enrichr performance evaluation, which demonstrates the success of the proposed method.

**Fig. 10.**
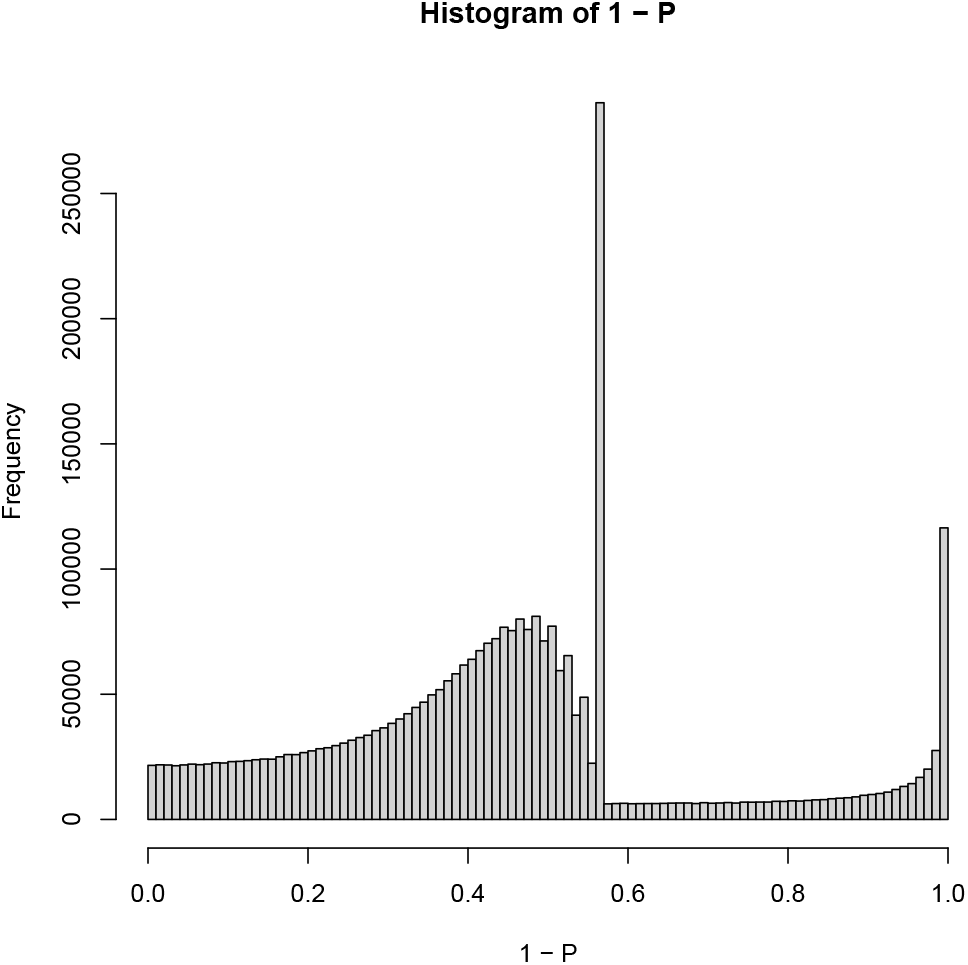
Histogram of 1 - *P_i_* computed by the proposed method using the GSE181596 dataset (H3K27me3)

### 3.3 TD-based unsupervised FE with optimized SD

All of the above experiments used PCA. To determine whether TD-based unsupervised FE with optimized SD worked as well, TD-based unsupervised FE with optimized SD was used to analyze the GSE74005 dataset. This dataset did not work well with the PCA-based unsupervised FE with optimized SD method. Histone modification was formatted as a tensor *x_ijk_* ∈ ℝ^*N*×16×2^ that represented H3K4me1 modifications in the *i*th region of the *j*th sample in the *k*th treatment (*k* =1: treated, *k* = 2: input). After obtaining SVVs, *u*_1*j*_ and *u*_2*k*_ were coincidences with the independence of the samples and the distinction between the treated and controlled samples. *G*(2,1, 2) had the largest absolute value, thus *u*_2*i*_ was used to select regions. As a result, a total of 70187 regions, the associated 14220 Entrez gene IDs, and 14187 gene symbols have been identified. Although these numbers are not so different from those in Table 3, performance is improved (Table 3). Thus, when PCA does not work, TD is worth testing as well.

## 4 Discussion

The proposed method has several advantages. First of all, it is a fully linear method that simply applies PCA or HOSVD to matrices and tensors. The only time-consuming process is SD optimization, which typically ends in a few minutes. The second advantage is that the proposed method accepts the bed or the bigWig file format. In contrast to bam/sam files that retain information about how individual short reads are mapped onto the genome, bed or bigWig files do not keep this information and instead retain which regions of the short read are mapped. The third advantage is robustness. Independent of samples, species, and platforms, the proposed method achieved similar performance levels. For example, the H3K27me3 modification is considered three times in Table 3; two are human ChIP-seq and one is mouse ChIP-seq. In spite of the different species, all three are associated with a similar number of gene symbols, 5208, 4994, and 3545, and they are all associated with the same number of enriched H3K27me3 profiles in the “Epigenomics Roadmap HM ChIP-seq” category of Enrichr. H3K27ac modifications were considered three times with two distinct protocols, ChIP-seq and CUT&Tag;. A similar number of gene symbols, 11590, 13061, and 15548 and the same number of H3K27ac-enriched profiles, 24, in the “Epigenomics Roadmap HM ChIP-seq” category of Enrichr were associated. Usually, although CUT&Tag requires protocols specific to CUT&Tag [19], the proposed method handled both ChIP-seq and CUT&Tag seamlessly. In addition to this, as seen in Tables 2 and 3, the proposed method successfully deals with all five “core histone marks” (H3K27ac, H3K4me3, H3K36me3, H3K27me3, and H3K9me3). Unfortunately, H3K4me1 cannot be properly dealt with using the proposed method. However, this is not a critical problem, since it can process H3K27ac modifications, which are strongly associated with H3K4me1 modifications [38]. Thus, the proposed method is still worth investigating.

In the above, *u_ℓi_* does not obey a Gaussian distribution, but instead fits a combination of two Gaussian distributions. To confirm these points, the analysis of the Gaussian mixture to *u*_1*i*_ of the GSE24850 dataset was applied (whose histogram 1 – *P_i_* is shown in Fig. 2). Then, the Bayesian information criterion of the Gaussian mixture drastically improves when the number of clusters increases from one to two and does not improve for more numbers of clusters. Table 4 shows the confusion matrix between clustering and selection of the proposed method when the number of assumed clusters varies from two to four. It is obvious that they are in agreement only when the number of clusters is assumed to be two as expected. All regions not selected by the proposed method are in one cluster and the majority of the selected regions belong to another cluster whereas this does not occur when the number of assumed clusters is either three or four. Thus, as expected, *u_ℓi_* did not obey a Gaussian distribution, but a mixture of two Gaussian distributions. Since the optimization of SD assumes a single Gaussian distribution, it is not expected to work well, but it does work well empirically (Table 3). The limitations of the proposed methods applied to histone modification must be investigated in the future.

**TABLE 4.** Confusion matrix between clustering by a mixture of Gaussian distributions(row) and selection by the proposed method (column)

|  |  | Number of assumed clusters |  |  |  |  |  |
| --- | --- | --- | --- | --- | --- | --- | --- |
|  |  | 2 |  | 3 |  | 4 |  |
|  |  | Proposed Method |  |  |  |  |  |
| Gaussian Mixture | Cluster | Adjusted $P$-value | | Adjusted $P$-value | | Adjusted $P$-value | |
|  |  | > 0.01 | ≤ 0.01 | > 0.01 | ≤ 0.01 | > 0.01 | ≤ 0.01 |
|  | 1 | 104902 | 428 | 99285 | 0 | 39555 | 0 |
|  | 2 | 0 | 874 | 5605 | 1222 | 61985 | 0 |
|  | 3 | — | — | 0 | 92 | 3350 | 1240 |
|  | 4 | — | — | — | — | 0 | 74 |

The proposed method has some weaknesses. If it is applied to non-core histone marks, it does not work well. For example, if it is applied to H3K9ac modifications, its performance is reduced. More detailed conditions by which the proposed method may be successfully applied to histone modification needs to be investigated.

In addition, we do not know how the proposed method outperforms the previous state-of-the-art methods. It is important to note that the proposed method is an unsupervised method. In contrast to the other methods that must assume some statistical properties on histone modification distribution along the genome, the proposed method assumes only that PCs and SVVs obtained obey Gaussian distributions. Other methods can be successfully applied to gene expression [13] and DNA methylation [14] because target specific methods can be developed. However, since no assumptions about the statistical properties of histone modification can be made, other methods are not robust when applied to identification of histone modification. The proposed method can infer histone modification correctly.

## 5 Conclusions

In this paper, a recently proposed PCA-based unsupervised FE with optimized SD was applied to identify histone modifications. It successfully counted five core histone marks and outperformed the state-of-the-art methods. The proposed method is expected to be a new state-of-the-art method for histone modification.

## Acknowledgments

This work was supported by KAKENHI [grant numbers 20H04848 and 20K12067] to YHT.

## Data Availability

Supplementary files and sample R code to perform analyses in this study can be found at https://github.com/tagtag/PCAUFEOPSDHM

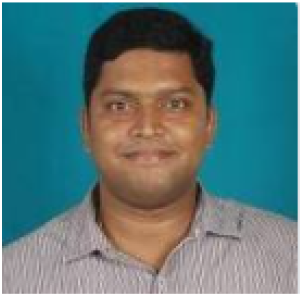

**Sanjiban Sekhar Roy** Dr. Sanjiban Sekhar Roy is an Associate Professor in the School of Computer Science and Engineering, Vellore Institute of Technology. He uses Deep Learning and machine learning techniques to solve many complex engineering problems, especially those are related to imagery. Dr Roy has vast experience in research specially in the field of advanced machine learning and deep learning. He is specialized in ML & deep convolutional neural networks towards solving various image related complex problem and various engineering problems such as computational biology, civil, energy inspired problems. Dr. Roy also has edited special issues for journals and many books with reputed international publishers such as Elsevier, Springer and IGI Global. Dr. Roy is adjunct researcher to Ton Duc Thang University, Vietnam since 2019.Very recently, Ministry of National Education, Romania in collaboration with “Aurel Vlaicu” University Arad Faculty of Engineers, Romania has awarded him with ‘Diploma of Excellence” as a sign of appreciation for the special achievements obtained in the scientific research activity in 2019.

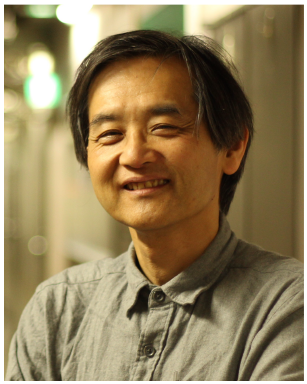

**Y-H. Taguchi** received a B.S. degree in physics from the Tokyo Institute of Technology and a Ph.D. degree in physics from the Tokyo Institute of Technology. He is currently a full professor with the Department of Physics, Chuo University, Japan. His works have been published in leading journals such as Physical Review Letters, Bioinformatics, and Scientific Reports. His research interests include bioinformatics, machine-learning, and non-linear physics. He is also an editorial board member of Frontiers in Genetics:RNA, PloS ONE, BMC Medical Genomics, Medicine (Lippincott Williams & Wilkins journal), BMC Research Notes, non-coding RNA (MDPI), and IPSJ Transaction on Bioinformatics.

